# A Poly(rC)-Binding Protein Mediates Iron Delivery to Ferritin in *Anopheles stephensi*

**DOI:** 10.64898/2026.08.04.742707

**Authors:** Chiging Tupe, Bharti Goyal, Tanwee Das De, Durva Bhatt, Bhuvan Dixit, Kailash C Pandey, Byoung-Kuk Na, S. Noushin Emami, Soumyananda Chakraborti

**Affiliations:** Birla Institute of Technology and Science Pilani, Hyderabad Campus, Hyderabad, India; ICMR-National Institute of Malaria Research, Dwarka, New Delhi, India; Academy of Scientific and Innovative Research (AcSIR), UP, India; Department of Medical Entomology and Zoology, ICMR-National Institute of Virology, Microbial Containment Complex, Pune, Maharashtra, India; Department of Parasitology and Tropical Medicine, Department of Convergence Medical Science, and Institute of Medical Science, Gyeongsang National University College of Medicine, Jinju 52727, South Korea; Natural Research Institute, University of Greenwich, Greenwich, UK; Department of Vector Biology, Liverpool School of Tropical Medicine, Liverpool, L3 5QA, UK; Molecular Attraction AB, Elektravägen 10, Hägersten, 126 30, Stockholm, Sweden; Department of Microbiology, Tumor and Cell Biology, Karolinska Institute, Stockholm, Sweden

**Author notes:** **Author Emails** Chiging Tupe, Bharti Goyal, Tanwee Das De, Durva Bhatt, Bhuvan Dixit, Kailash C Pandey, Byoung-Kuk Na, S. Noushin Emami, Soumyananda Chakraborti.

**Keywords:** Iron homeostasis, Poly(rC)-binding protein (PCBP), Iron chaperone, host-pathogen interaction, Malaria

## Abstract

Iron homeostasis is essential for both the *Plasmodium* parasite and its hosts, the mosquito and human; however, the molecular mechanisms governing intracellular iron trafficking remain poorly understood. In mammals, poly(rC)-binding proteins (PCBPs) function as major iron chaperones by delivering Fe^2+^ to ferritin and other iron-dependent proteins, yet their role in insects has not been investigated. Here, we provide the first structural and functional characterization of the PCBP from a major malaria vector, *Anopheles stephensi*. Comparative structural and evolutionary analyses showed that insect PCBPs are most closely related to human PCBP3. Coordinated expression and co-localization of *As*PCBP with ferritin following blood feeding suggested a conserved role in mosquito iron homeostasis. Functional analyses demonstrated that KH2-KH3 linker and KH3 domain of PCBP, plays an important role in the *As*PCBP-ferritin interaction and that deletion of these regions, unexpectedly increased ferritin binding affinity but impaired iron delivery, indicating that these regions are dispensable for complex formation but essential for efficient iron transfer. Collectively, our results identify PCBP as a key component of mosquito iron homeostasis that could be exploited for the development of novel antimalarial strategies.

## Introduction

Iron metabolism is a fundamental biological process in all organisms and plays a critical role in disease pathogenesis, including malaria. Hematophagous insects such as mosquitoes and ticks largely depend on iron for their growth and reproductive fitness (1). Similarly, *Plasmodium* parasites also require iron for their growth and replication within host cells, making iron homeostasis a critical determinant of both vector and parasite biology (2). Previous studies have shown that iron deficiency can confer protection against *Plasmodium* infection in both mosquito and human host (3–5). However, despite this growing evidence, the exact mechanisms of iron metabolism remain largely unexplored specially in the context of malaria pathogenesis and eradication efforts.

There are several key proteins involved in iron metabolism that play important roles in the host-parasite interplay. For instance, ferritin whose function as an intracellular iron storage protein is conserved across species, limits the availability of free iron to invading pathogens (6). Similarly, the same study demonstrated that the divalent metal transporter 1 (DMT1) localized to the parasite digestive vacuole plays a crucial role in iron distribution (6, 7). Another host protein, hepcidin, although not reported in insects, is a key regulator of systemic iron homeostasis in vertebrates, controlling circulating iron levels by inhibiting the iron exporter ferroportin, thereby reducing the availability of serum iron (8). In addition to these well-established regulators, a multifunctional iron- and RNA/ssDNA-binding protein(9–12), poly(rC)-binding protein (PCBP), that functions as a major intracellular iron chaperone in mammals, has recently been identified as a target of parasites such as *Leishmania* for host iron acquisition. (13).

PCBPs belong to the family of heterogeneous nuclear ribonucleoproteins (hnRNPs) and are represented by four isoforms in humans (PCBP1-4)(14). Each isoform contains three highly conserved K-homology (KH) domains that harbor both iron- and RNA/ssDNA-binding residues (two residues per domain), structural variation among PCBPs primarily arises from differences in the KH2-KH3 linker region (15). Different PCBP isoforms are known to interact with different proteins for iron delivery (16–19). All isoforms can bind up to three ferrous iron (Fe^2+^) ions (20) and facilitate their safe intracellular delivery to target proteins such as ferritin, heme oxygenase-1 (HO-1), DMT1,etc (14, 17, 18, 21). Additionally, PCBPs have been reported to play an essential role in erythrocyte maturation, with PCBP1 deficiency in mice resulting in microcytic anaemia (22). In addition to their well-established role as post-transcriptional regulators, PCBPs also function as iron chaperones and play critical roles in cellular iron homeostasis and haematopoiesis (22, 23). Notably, PCBP1 deprivation in PCBP1-deficient mice resulted in microcytic anaemia (22). Among all the proteins that PCBP delivers iron to, ferritin has been identified as a major iron-accepting partner, facilitating the safe storage of intracellular iron(14).

Pathogens have evolved strategies to disrupt this PCBP-mediated iron delivery pathway. A recent and notable example is observed in *Leishmania*, where PCBP1/2 are cleaved at the KH2–KH3 linker region by a metalloprotease, GP63, thereby impairing its ability to deliver iron to ferritin and increasing the intracellular labile iron pool available for parasite survival(13). Similarly, viruses such as poliovirus and hepatitis virus cleave host PCBP at a comparable site using the viral 3C protease (a cysteine protease), impairing its role in host translation and promoting viral replication; knockdown of PCBP resulted in a decrease in viral yield. (24–26). In addition to infectious diseases, PCBPs have also been implicated in cancer, where they play an important function in tumor progression (27–29). Together, these findings highlight the critical role of PCBPs in host iron homeostasis and underscore their potential exploitation by diverse pathogens.

PCBPs in mammals, particularly PCBP1 and PCBP2, have been extensively studied with respect to their structural characterization, biological functions, and roles in disease. However, information on PCBPs in other phyla, as well as their evolutionary origins remains limited. In insects, only the PCBP homolog in *Drosophila*, annotated as mushroom body expressed (mub), has been functionally characterized, where it has been implicated in the regulation of circadian rhythm, similar to the mammalian PCBP1, by enhancing the interaction of circadian repressor genes such as CRY1 with the CLOCK/BMAL1 complex.(30). Given that insect vectors such as mosquitoes play a critical role in disease transmission, greater attention to this important iron chaperone is warranted.

This study provides the first structural and functional characterization of an insect poly(rC)-binding protein (PCBP) from the malaria vector *Anopheles stephensi*. The mosquito PCBP was found to be more closely related to human PCBP3, while retaining conserved RNA/ssDNA-binding residues but exhibiting substitutions in key iron-binding residues within the KH1 and KH3 domains. *As*PCBP and ferritin showed coordinated expression and co-localization following a blood meal, supporting their role in mosquito iron homeostasis. The KH3 domain together with the KH2-KH3 linker was found to be essential for efficient iron transfer to ferritin, despite deletion of these regions increasing ferritin-binding affinity. Overall, these findings identify *As*PCBP as a one of the key regulators of iron trafficking in mosquitoes.

## Results

### Divergence of Iron-Binding Residues in PCBPs across evolution

In humans, PCBP exists as four isoforms; however, unlike their mammalian counterparts, only a single PCBP isoform has been reported in insects according to VectorBase. PCBPs consist of three K-homology (KH) domains (Figure 1A-C) and all three KH homology domains are structurally highly conserved across isoforms in humans and even when compared with insect PCBP, as seen in *Anopheles stephensi* PCBP (RMSD < 1 Å) (Figure 1D), with major structural differences arising from the KH2-KH3 linker region (Figure 1A, Table 1). Comparison of the full-length protein sequences revealed that *Anopheles stephensi* PCBP (*As*PCBP) most closely resembles human PCBP3 (~54%) (Table 1), an isoform that remains relatively understudied compared with the other mammalian PCBPs. However, despite the conservation of RNA-binding sites and physicochemical properties across species (Table 2), insect PCBPs appear to differ in their iron-binding residues (Figure 1C), which have been reported to be essential in human PCBP isoforms(17, 24–26).

**Table 1.** Sequence-based percentage identity of mosquito PCBP with its mammalian homologs.**pecies Protein**.

| Species | Protein | Sequence similarity to *AsPCBP* |  |  |  |  |
| --- | --- | --- | --- | --- | --- | --- |
|  |  | Full protein | KH1 | KH2 | KH3 | KH2-KH3 Linker |
| *Anopheles stephensi* | *AsPCBP*<br>(A0A182YKR7) | 100% | 100% | 100% | 100% | 100% |
| *Anopheles gambiae* | *AgPCBP*<br>(A0A903X4J2) | 94.59% | 100% | 100% | 100% | 58.97% |
| *Aedes Aegypti* | *AaPCBP*<br>(A0A6I8T4P0) | 80.58% | 98.63% | 98.65% | 100% | 57.26% |
| *Drosophila melanogaster* | *DmPCBP*<br>(M9NEH9) | 69.11% | 90.28% | 91.89% | 83.33% | 46.43% |
| *Homo sapiens* | *HsPCBP1*<br>(Q15365) | 46.69% | 66.67% | 63.01% | 53.25% | 24.55% |
| *Homo sapiens* | *HsPCBP2*<br>(Q15366) | 49.03% | 66.67% | 64.38% | **58.67%** | 32.17% |
| *Homo sapiens* | *HsPCBP3*<br>(P57721) | **53.98%** | **68.06%** | **73.24%** | 55.84% | **40.86%** |
| *Homo sapiens* | *HsPCBP4*<br>(P57723) | 45.17% | 58.33% | 65.71% | 55.44% | 27.94% |

**Table 2.** Physicochemical properties of PCBP homologs across different species, calculated using ProtParam.

| Species | No.of amino acids | Molecular weight | Isoelectric point | Hydrophobicity (GRAVY)* | Dominant amino acids |
| --- | --- | --- | --- | --- | --- |
| *Anopheles stephensi* | 380 | 41.8 kDa | 6.98 | -0.061 | His(H), Gly (G), Ala (A) |
| *Anopheles gambiae* | 391 | 43 kDa | 8.43 | -0.059 | Ile(I), Gly (G), Ala (A) |
| *Aedes Aegypti* | 365 | 38.5 kDa | 8.27 | -0.142 | Ile(I), Ala (A), Gly (G) |
| *Drosophila melanogaster* | 386 | 40.7 kDa | 8.53 | -0.109 | Ile(I), Ala (A), Gly (G) |
| *Homo sapiens*<br>(*HsPCBP1*) | 356 | 37.5 kDa | 6.66 | -0.106 | Gly (G), Ser (S), Ile (I) |
| *Homo sapiens*<br>(*HsPCBP2*) | 364 | 38.5 kDa | 6.33 | -0.135 | Ser (S), Gly (G)/Ile (I), Ala (A) |
| *Homo sapiens*<br>(*HsPCBP3*) | 371 | 39.5 kDa | 8.22 | -0.206 | Gly (G), Ile (I), Ser (S) |
| *Homo sapiens*<br>(*HsPCBP4*) | 403 | 41.3 kDa | 8.42 | 0.065 | Ala (A), Gly (G)/Ser (S), Pro (P) |

\*Negative values denote a hydrophilic nature

**Figure 1.**
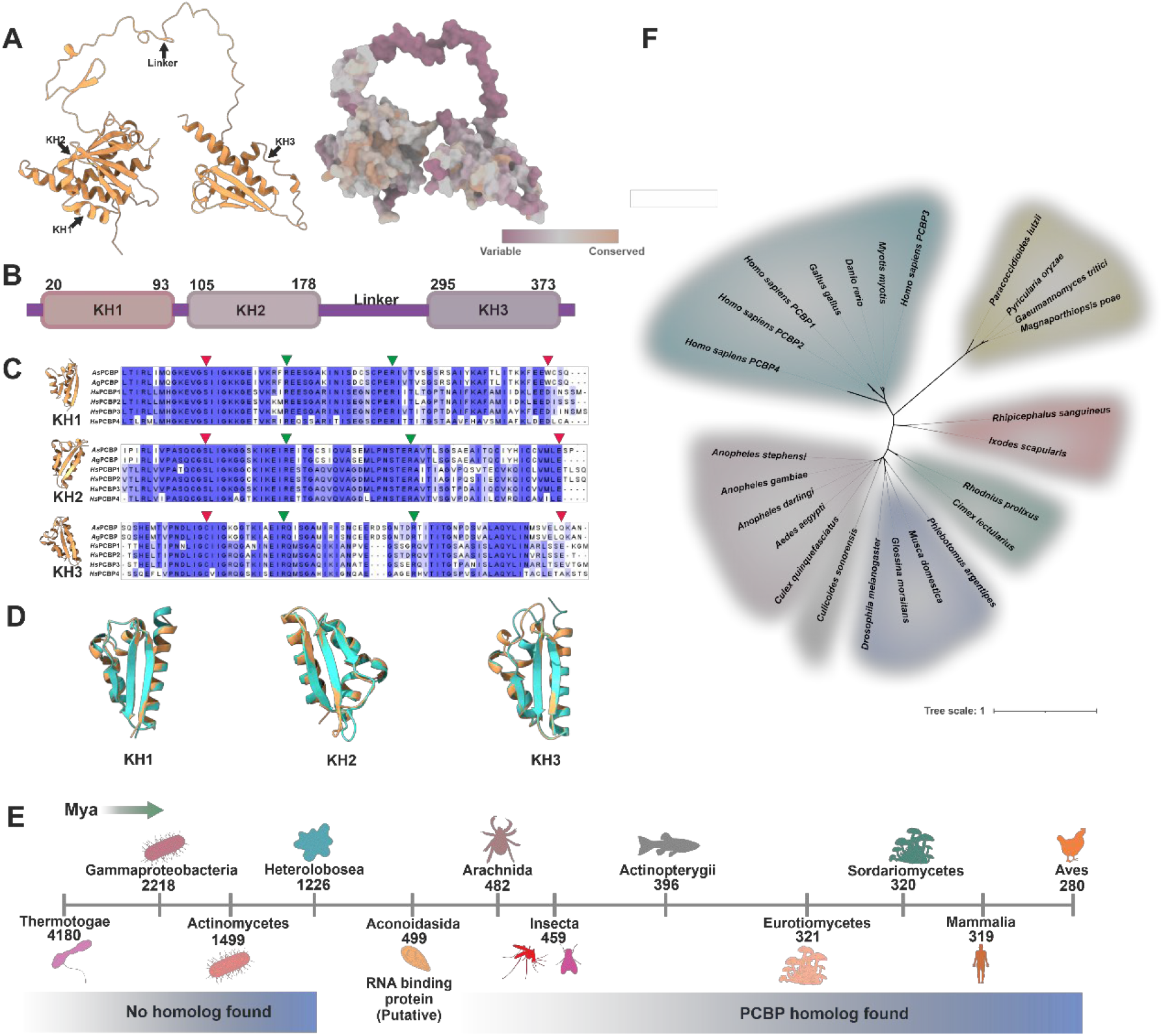
Structural, evolutionary, and phylogenetic analysis of poly(rC)-binding protein (PCBP). (A) Predicted structure of *Anopheles stephensi* PCBP (*As*PCBP) shown as ribbon (left). Residue conservation (right) across different species was mapped onto thestructure usingthe AL2CO algorithm in ChimeraX, whereconserved residues are shown in orange and variable residues in magenta.(B) domain diagram of *As*PCBP showingthe three K-homology (KH) domains, connected by a 117 amino acid long variable linker region (KH2-KH3 linker). (C)Multiple sequence alignment of the KH domains of *As*PCBP with *Ag*PCBP (*Anopheles gambiae*) and the four human homologs (*Hs*PCBP1-4). Conserved residues involved in iron (red arrow) and RNA binding (green arrow) are highlighted, showing strong conservation of key functional motifs across species. (D) Superposition of the three KH domains of *As*PCBP (orange) with the corresponding domains of *Hs*PCBP3 (cyan), demonstrating structural conservation of the KH domains (RMSD <1Å). (E) Evolutionary distribution of different classes of organisms mapped onto a divergence timeline (million years ago, Mya) using divergence estimates from TimeTree (https://timetree.org/). The presence or absence of PCBP homologs in each lineage is indicated. (F) Maximum-likelihood phylogenetic tree of PCBP homologs identified across representative taxa. The tree was constructed with using Le Gascuel substitution model, 1,000 bootstrap replicates, illustrating the evolutionary relationships among PCBP homologs from diverse lineages.

For instance, in *Anopheles stephensi*, iron-binding residues D114 (KH1 domain) and E338 (KH3 domain) of human PCBP3 are replaced by tryptophan W90 (KH1 domain) and glutamine Q370 (KH3 domain), respectively (Figure 1B). Since PCBPs bind three Fe^2+^ atoms through coordination with iron-binding residues distributed across all three KH domains (two residues per domain), as demonstrated in a previous study in which mutations in these residues significantly reduced the iron-binding affinity of human PCBP1 (15), these variations may potentially affect their iron-binding capability. It is important to note that the iron-binding residues, and consequently the iron-binding site, may also differ from those of the human homolog. Therefore, we predicted the potential iron-binding sites using the MIB2 server (31) as well (Table S1). That being said, the iron-binding residues of PCBP are hypothesized to have evolved more recently than the evolutionarily conserved RNA-binding residues found in KH domain-containing proteins such as hnRNPK (32). Consistent with this hypothesis, iron-binding residues are absent in the KH domain containing proteins of the prokaryotes as well as in evolutionarily older eukaryotic lineages (Figure 1E). Phylogenetic analysis further revealed that, among the four human PCBP isoforms, PCBP3 forms a distinct clade (Figure 1F). Its closer relationship to homologs from evolutionarily older taxa, together with its comparatively lower iron chaperone activity than the other PCBP isoforms (14), suggests that PCBP3 may represent a more ancestral form of the protein.

Consecutively, we performed domain-specific surface electrostatic potential mapping and found that, the canonical iron-binding sites seen in human and insect PCBPs, are rich in negatively charged residues. While the RNA binding sites had more positively charged residues, such as arginine and lysine, which have a higher affinity for RNA (Table S2).

### *As*PCBP and Ferritin Exhibit Coordinated Expression Post Blood Meal

In mammals, PCBPs function as primary iron chaperones for ferritin (14), delivering Fe^2+^ to the latter for storage (14). Therefore, a correlation between their expression and cellular localization is expected. To validate this, we employed multiple approaches. First, we performed immunofluorescence assays to examine the co-localization of *As*PCBP and ferritin in the whole body, midgut, and ovaries of blood-fed adult female *Anopheles stephensi* mosquitoes at different post-blood meal time points.

As the genomic database for *Aedes aegypti* are more comprehensive and better annotated than those for *Anopheles stephensi*, we performed relative gene expression analyses of PCBP, ferritin, and frataxin, homolog of the major mitochondrial iron chaperone in mammals, across different developmental stages and tissues of *Aedes aegypti* to complement the transcriptomic data available in VectorBase. It is noteworthy that these genes exhibit high structural and sequence conservation across mosquito vectors (Table 1) (33), supporting the relevance of using *Aedes aegypti* as a comparative model instead.

The results revealed that the expression patterns of ferritin and *As*PCBP were highly similar, with peak expression observed at 48-72 h post-blood meal in the abdominal region, where the midgut and ovaries are located (Figure S1A), as well as in the developing reproductive tissues of third-instar *Anopheles stephensi* larvae (Figure S1A). A similar expression pattern was observed in the midgut and ovaries of adult female *Anopheles stephensi* and was further corroborated by the qPCR results obtained from *Aedes aegypti* (Figure 2A-E), consistent with the ferritin expression profile reported in previous studies (33–35). These findings suggest that the increased demand for ferritin to cope with the influx of iron during blood digestion is accompanied by the upregulation of its iron chaperone, *As*PCBP, thereby facilitating efficient iron delivery to ferritin for storage and minimizing reactive oxygen species (ROS)-mediated damage.

**Figure 2.**
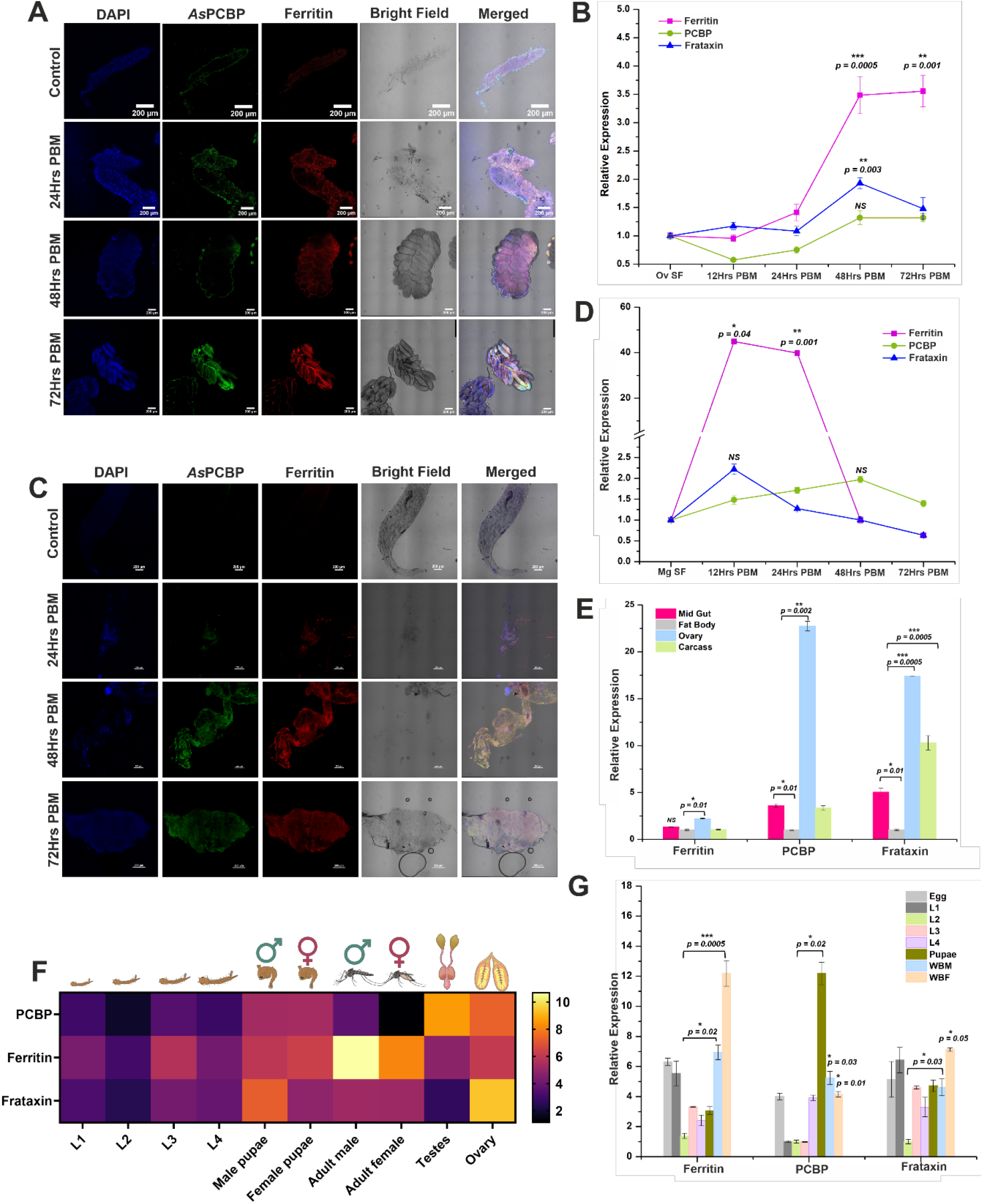
Comparative localization and expression profiling of iron metabolism-associated proteins in *Anopheles stephensi*and *Aedes aegypti*. (A & C) Immunofluorescencelocalizationof *As*PCBP (Alexa488, green) and ferritin (Alexa647, red) in the (A) ovaries and (C) midguts of female *Anopheles stephensi* mosquitoes at 24, 48, and 72 h post-blood meal (PBM), with sugar-fed (SF) females serving as controls. Nuclei were stained with DAPI (blue). Representative bright-fieldand merged images are shown. Scale bars = 200 μm, Magnification-20X. (B) Relative transcript expression of ferritin, PCBP, and frataxin in the (B) ovaries and (D) midgut of female *Aedes aegypti* mosquitoes at 12, 24, 48, and 72 h PBM relative to sugar-fed (SF) controls. (E) Tissue-specific expression profile of ferritin, PCBP, and frataxin in sugar-fed female *Aedes aegypti* mosquitoes. Transcript levels were measured in the midgut, fat body, ovary, and carcass using qPCR. (F) Heatmap showing developmental stage-specific transcript abundance of ferritin, PCBP, and frataxin in *Aedes aegypti* obtained from the available transcriptomic datasets of VectorBase, Transcripts Per Million (TPM) values were normalized for relative expression visualization, raw data is provided in the SI. (G) Relative transcript expression of ferritin, PCBP, and frataxin across different developmental stages of *Aedes aegypti*, including eggs, 1^st^ to 4^th^ instar larvae (L1sL4), pupae, whole-body males (WBM), and whole-body females (WBF). Data in panels B, D, E, and G are presented as mean ± SEM from three independent biological replicates (N=3). Statistical significance was determined using Student’s t-test. Significance levels are denoted as p^***^ ≤ 0.0005, p ^**^≤ 0.001,0.002, p^*^ ≤ 0.01,0.02,0.03 and 0.05; NS= not significant.

To gain a more comprehensive understanding of iron chaperoning, we simultaneously examined the expression of frataxin, which, unlike PCBP, is not a known interacting partner of ferritin. Interestingly, the relative expression of frataxin was higher than that of PCBP following blood feeding (Figure 2B&D). As a key component of the mitochondrial iron-sulfur (Fe-S) cluster assembly machinery, frataxin facilitates the biogenesis of Fe-S proteins, including iron regulatory protein 1 (IRP1) (36). Given the central role of IRPs in regulating ferritin expression in response to intracellular iron levels, the elevated expression of frataxin may reflect the increased requirement for coordinated iron homeostasis during blood digestion. In the development stages of *Aedes aegypti*, PCBP and ferritin exhibited similar expression patterns (Figure 2E&F), with the highest expression observed in third-instar larvae, pupae, and adult males, whereas frataxin expression peaked during the pupal stage (Figure 2E).

In mammals, PCBP1 and PCBP2 serve as the primary cytosolic iron chaperones, and their knockout resulted in severe developmental abnormalities and embryonic lethality in mice (37). Therefore, we attempted performing RNAi-mediated knockdown of PCBP in *Aedes aegypti*. Unfortunately, despite repeated efforts to optimize the RNAi conditions, we were unsuccessful. Therefore, the precise physiological role of PCBP in mosquito iron metabolism remains to be elucidated and warrants further investigation.

### Comparative Ferritin Binding of PCBP Constructs

Previous studies have shown that PCBP can directly bind to ferritin, either alone or in complex with glutathione (GSH), to deliver iron (14, 16). However, the exact coordinates of this interaction remain unclear. Based on docking analysis, we found that in humans, *Hs*PCBP1 and *Hs*PCBP2, which are reported to be the primary iron chaperones among the four isoforms(14), preferentially bind to the threefold pore channel of the ferritin cage (Figure S2A-B). These observations are consistent with the threefold pore being the major route for iron influx into ferritin for storage(38). In contrast, docking of *Hs*PCBP3 with the human ferritin cage(Hftn) revealed binding at the interface between the three-fold and four-fold channels (Figure S2B). A similar binding pattern was also observed for *As*PCBP with mosquito ferritin (MqFtn) of *Anopheles stephensi* (Figure S2D). Docking analysis further suggested that the KH2-KH3 linker region contributes significantly to this interaction across PCBPs (Figure S2A-E). Therefore, a truncated construct lacking the KH2-KH3 linker and KH3 domain (*As*PCBPΔKH3) was generated, cloned, and purified (Figure 3A&B).

**Figure 3.**
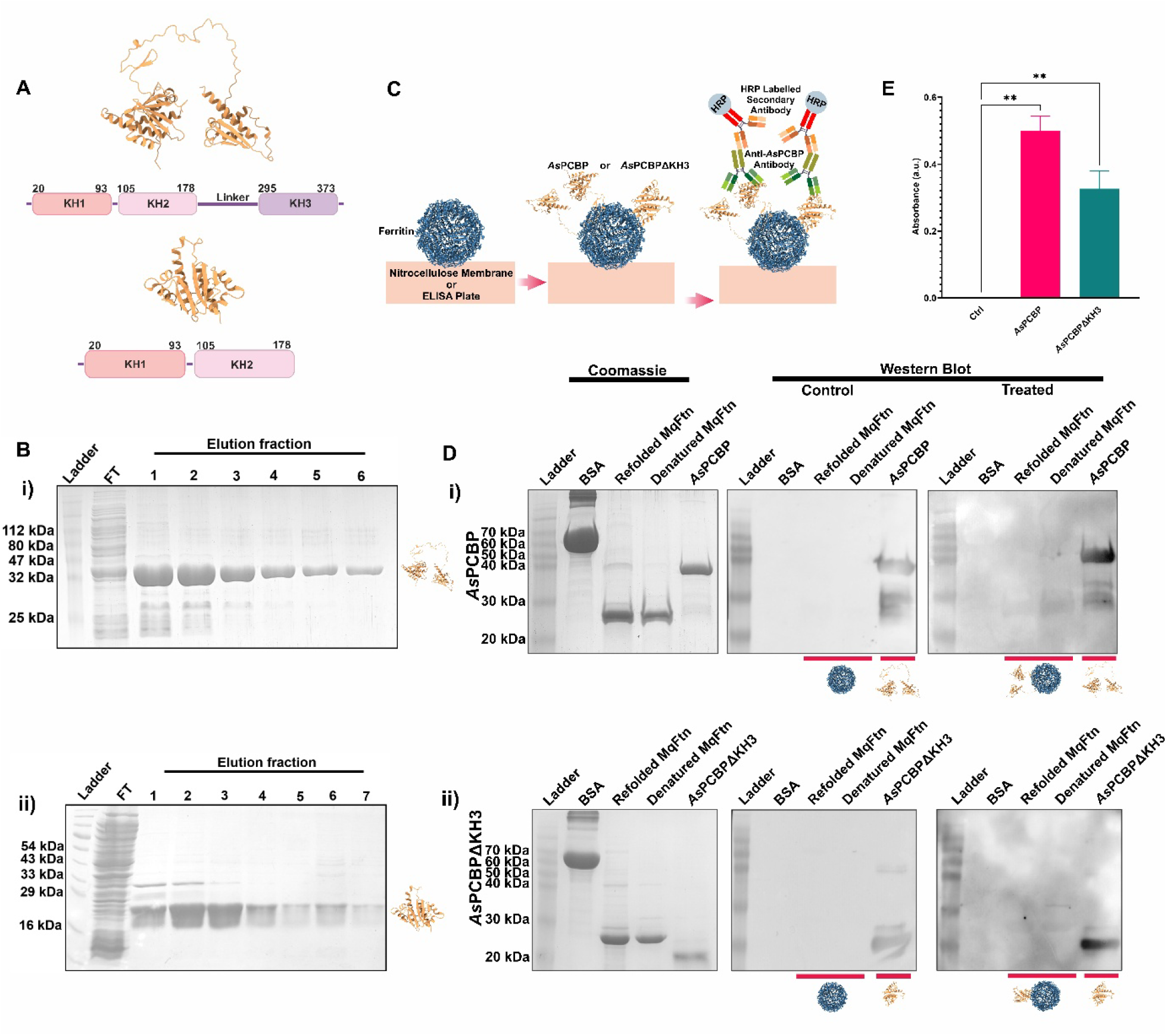
Protein-protein interaction between *As*PCBP (full-length) and *As*PCBPΔKH3 (Truncated KH3 domain) and MqFtn. (A) Structural and domain diagram of *As*PCBP and *As*PCBPΔKH3 (B) SDS PAGE gel image showing purified (i) *As*PCBP and (ii) *As*PCBPΔKH3 (C) Schematic representation of the Far-Western and ELISA assay performed.(D) Far-Westernblot analysis demonstratinginteraction between MqFtn and(i) *As*PCBP or (ii) *As*PCBPΔKH3. MqFtn (prey) was immobilized on a nitrocellulose membrane, followed by incubation with recombinant *As*PCBP or *As*PCBPΔKH3 (bait). BSA was used as a negative control. Bound PCBP was detected using anti-AsPCBP antibody. Specific signal at the position corresponding to MqFtn indicates protein-protein interaction. Coomassie-stained gels are shown alongside the corresponding control and treated immunoblots. (E) Quantification by ELISA performed under conditions similar to the Far-Western assay, showing binding of *As*PCBP and *As*PCBPΔKH3 to MqFtn. Data are presented as mean ± SD from three independent biological replicates (^**^p < 0.005, one-way ANOVA).

To experimentally validate the interaction between *As*PCBP and MqFtn and to evaluate the effect of KH3 domain truncation, Far-Western blotting and ELISA were performed. In these assays, MqFtn (bait) was immobilized on nitrocellulose membranes or ELISA plates and incubated with either full-length *As*PCBP or *As*PCBPΔKH3 (prey) (Figure 3C). Immunodetection using polyclonal anti-*As*PCBP antibodies revealed signals corresponding to the position of MqFtn, confirming the formation of both the *As*PCBP-MqFtn and *As*PCBPΔKH3-MqFtn complexes, as the antibodies specifically detected the prey proteins bound to immobilized MqFtn (Figure 3C&D).

Interestingly, *As*PCBPΔKH3 retained its ability to bind MqFtn (Figure 3C&D), although a modest reduction in band intensity and ELISA absorbance was observed compared with the full-length protein (Figure 3D&E). Since the anti-*As*PCBP antibody was polyclonal and was raised against the full-length *As*PCBP protein, the truncated construct lacks a proportion of the antibody-recognized epitopes. Therefore, the reduced immunoreactivity is likely attributable to decreased antibody recognition rather than a substantial reduction in the interaction between *As*PCBPΔKH3 and MqFtn. Additionally, the electrostatic surface potential maps of these docked structures revealed that the interaction between PCBPs and ferritin is predominantly electrostatic, with increase in positively charged residues interactions upon truncation (*As*PCBPΔKH3) (Figure S3). To further assess their interaction, we performed microscale thermophoresis (MST) to quantify the binding affinity of *As*PCBP and *As*PCBPΔKH3 toward MqFtn. MST revealed a lower binding affinity for full-length *As*PCBP compared with *As*PCBPΔKH3 (K_d_ = 51.3 ± 14.56 µM vs. 26.65 ± 14.2 µM) (Figure 4A-B). A similar trend was observed for binding to human ferritin (Hftn), with K_d_ values of 2.45 ± 0.13 µM for *As*PCBP and 0.26 ± 0.01 µM for *As*PCBPΔKH3 (Figure S4A-B).

**Figure 4.**
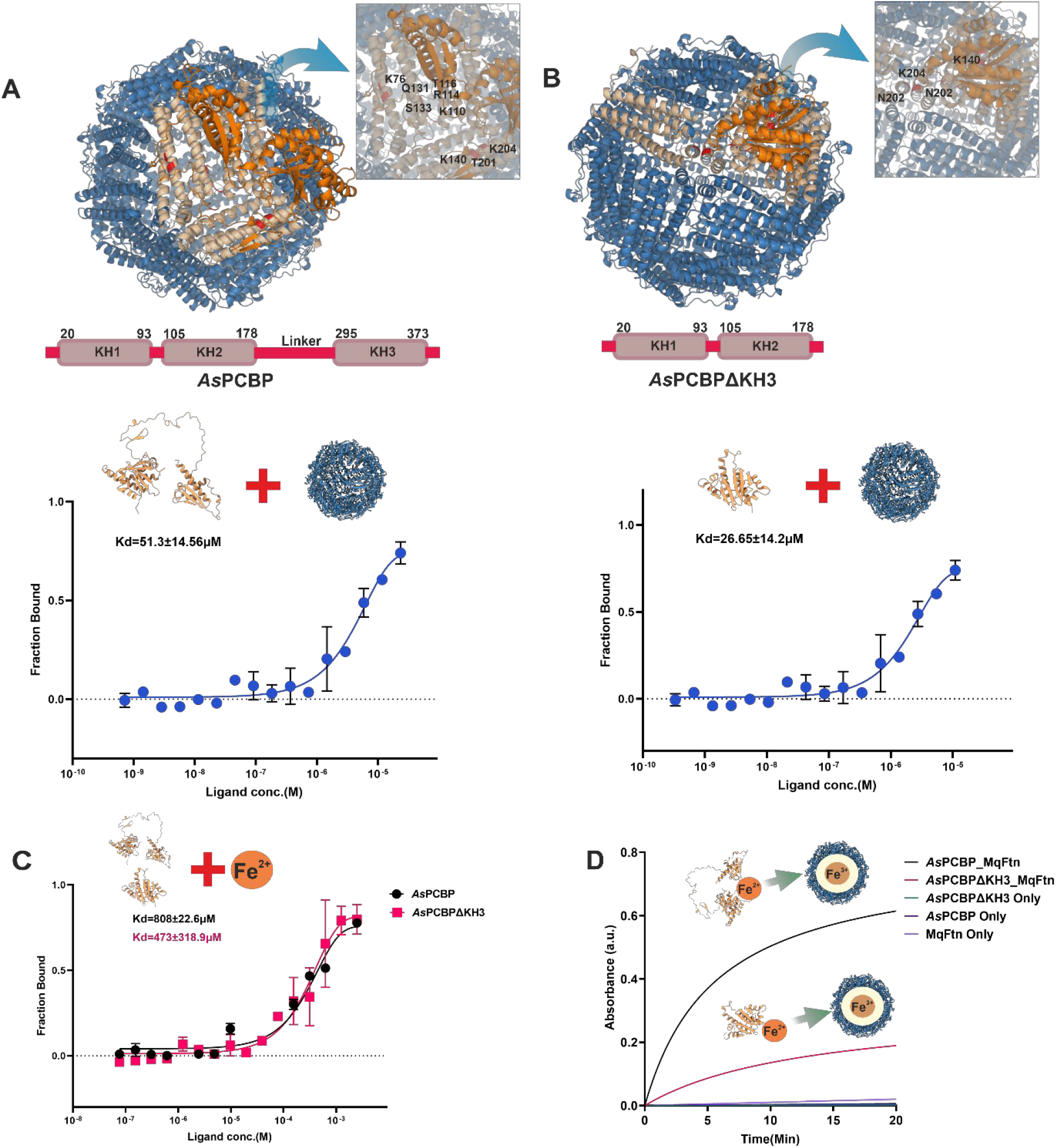
Structural and functional analysis of MqFtn interaction with *As*PCBP and *As*PCBPΔKH3. (A-B) Structural models of docked complexes and the microscale thermophoresis (MST) analysis showing binding of between MqFtn and (A) full-length *As*PCBP or (B) *As*PCBPΔKH3; figures were generated using pymol and dockings were performed using ClusPro 2.0. Insets highlight interacting residues on MqFtn. The domain diagram of *As*PCBP and *As*PCBPΔKH3 is shown below for reference. For MST,data represent duplicate measurements plotted as fraction bound versus ligand concentration, with calculated K_d_ values indicated, (C) MST-based binding analysis of *As*PCBP and *As*PCBPΔKH3 with Fe^2+^ (FeSO_4_), (D) Iron oxidation assay measuring iron loading into MqFtn in the presence of *As*PCBP or *As*PCBPΔKH3 at 310nm. Controls include *As*PCBP alone, *As*PCBPΔKH3 alone, and MqFtn alone. Increased absorbance indicates Fe^2+^ to Fe^3+^ oxidation. Truncation of *As*PCBP increased its binding affinity to ferritin but decreased its iron delivering efficiency to MqFtn.

Finally, the binding affinityof ferrous iron (Fe^2+^, supplied as FeSO_4_) to both constructs was evaluated. Consistent with the above observations, FeSO_4_ exhibited a lower binding affinity for full-length *As*PCBP compared with *As*PCBPΔKH3 (K_d_ = 808 ± 22.6 µM vs. 473 ± 318.9 µM) (Figure 4C).

### Effect of KH3 Deletion on Iron Delivery to Ferritin

The KH domains of PCBPs play a crucial role in their activity and interactions with various proteins. For instance, the KH2 domain of PCBP2 interacts with the N-terminal region of DMT1 to receive imported Fe^2+^ and with the C-terminal region of ferroportin to donate Fe^2+^ for export (39). Similarly, heme oxygenase 1 (HO-1) interacts with the KH3 domain of PCBP2 to facilitate the transfer of Fe^2+^ released during heme degradation (17); therefore, we examined the effect of domain deletion on their physiological activity. To examine this, an iron oxidation assay was performed using *As*PCBP or *As*PCBPΔKH3 in combination with MqFtn to assess their Fe^2+^ delivery capacity to MqFtn. In contrast to the binding affinity results, deletion of the KH3 domain reduced iron delivery to ferritin, as reflected in K_m_ (13.23 ± 0.06 µM for *As*PCBPΔKH3 compared with 5.657 ± 0.26 µM for full-length *As*PCBP) (Figure 4D), suggesting that higher binding affinity is not directly proportional to higher activity.

To further evaluate whether iron loading or depletion of *As*PCBP or *As*PCBPΔKH3 affects the reaction kinetics, the proteins were preincubated with 100 µM FeSO_4_ or EDTA, followed by extensive dialysis. It is important to note that higher concentrations of either treatment caused drastic protein precipitation. Interestingly, neither treatment resulted in a significant change in the kinetic parameters (Figure S4C), suggesting that the Fe^2+^ sequestered by PCBPs from the LB medium during culturing was very strongly coordinated and could not be effectively chelated by EDTA. This may also explain the lower binding affinity for FeSO_4_ observed in the MST results above.

## Discussion

In pathogenesis and disease transmission, the acquisition of nutrients from the host through parasitism plays an indispensable role for both the parasite and the vector; therefore, understanding these interactions is critical for developing effective strategies for disease control and eradication. Iron homeostasis represents a central component of the host-parasite interface in malaria (2, 5), yet the molecular mechanisms governing iron trafficking between mosquito/human, and *Plasmodium* remain incompletely understood.

The higher similarity of insect PCBP homologs with human PCBP3 raises the possibility that this isoform more closely resembles the ancestral PCBP from which the vertebrate isoforms diversified. A similar evolutionary scenario has previously been proposed for human PCBP1, which is thought to have arisen more recently through retrotransposition of PCBP2(37). Accordingly, it is plausible that gene duplication followed by functional specialization gave rise to the highly efficient iron chaperones PCBP1 and PCBP2 in vertebrates, whereas insects retained a single multifunctional homolog capable of coordinating both RNA metabolism and intracellular iron trafficking. Although this hypothesis remains speculative, it provides a plausible evolutionary explanation for the presence of only a single annotated PCBP homolog in mosquitoes despite the substantial iron demands associated with hematophagy.

Additionally, when compared to human PCBP, mosquito PCBP displayed high conservation of KH-domain architecture characteristic of the hnRNP family. Despite this structural conservation, two of the six reported iron-binding residues present in mammalian PCBPs were substituted in the mosquito homolog (Figure 1C). Previous mutagenesis studies have demonstrated that these residues are essential for Fe^2+^ coordination in human PCBPs (15), suggesting that the mosquito protein may possess distinct iron-binding properties or employ an alternative mode of metal coordination. Interestingly, the RNA-binding residues remained almost completely conserved across the analysed taxa (Figure 1C), supporting the hypothesis that the RNA-binding function of KH domains predates the acquisition of iron-chaperone activity (32). Our phylogenetic analyses further support this evolutionary scenario, demonstrating that the appearance of canonical iron-binding residues coincides with more recently diverged eukaryotic lineages, whereas these residues are absent from prokaryotes and evolutionarily older organisms (Figure 1E&F). Although experimental validation of this evolutionary model will require structural determination of iron-bound insect PCBPs using approaches such as cryo-EM or X-ray crystallography would be required to resolve these questions, our findings suggest that iron chaperone activity may have evolved through functional adaptation of an ancestral RNA-binding protein rather than through the emergence of an entirely new protein family.

The coordinated expression of PCBP and ferritin following blood feeding further supports a conserved role for PCBP in mosquito iron homeostasis (Figure 2A-G). Both proteins exhibited similar temporal and spatial expression profiles, with maximal abundance in the midgut and ovaries during active blood digestion, consistent with the period of highest iron influx. This coordinated expression suggests that intracellular iron trafficking and ferritin-mediated storage are tightly coupled processes that function to minimize the accumulation of cytosolic labile iron while ensuring sufficient iron availability for oogenesis. However, it is important to note that PCBP also plays a major role in the post-transcriptional regulation of numerous genes following blood feeding (11, 37, 43). Therefore, the observed increase in its expression may also reflect its involvement in these regulatory processes. The elevated expression of frataxin further indicates that mitochondrial iron utilization is simultaneously enhanced, highlighting the coordinated regulation of multiple iron-handling pathways following blood feeding.

A major mechanistic finding of this study is the identification of the KH2-KH3 linker and KH3 domain as important regulators of ferritin-mediated iron delivery (Figure S2A-E). Previous studies have shown that different KH domains mediate interactions with proteins involved in iron metabolism (16–18); however, the regions of PCBP responsible for ferritin binding have remained unclear. Surprisingly, deletion of the KH2-KH3 linker together with the KH3 domain increased ferritin binding affinity while significantly impairing iron delivery (Figure 3A-D). These findings suggest that strong protein-protein binding alone is not sufficient for efficient iron transfer and that the KH2-KH3 linker facilitates this process. We therefore propose that the KH2-KH3 linker is probably acting as a flexible regulatory region that facilitates efficient iron transfer from PCBP to ferritin. High-resolution structural studies of ferritin-PCBP complexes will be necessary to define the molecular basis of this mechanism and to determine how linker flexibility contributes to iron trafficking.

To summarize, this study provides the first characterization of an insect poly(rC)-binding protein (*As*PCBP) from the malaria vector *Anopheles stephensi*, along with a comparative and functional analysis of its interaction with mosquito ferritin following a blood meal. We also present quantitative evidence on how KH3 domain truncation affects the *As*PCBP-MqFtn interaction and iron delivery.

It is important to acknowledge the limitations of this study. In particular, high-resolution structural studies of the PCBP-ferritin complex will be required to define the molecular mechanism underlying PCBP-mediated iron transfer. Future studies should identify additional proteins involved in the ferritin-PCBP axis and resolve the structure of the PCBP-ferritin complex using high-resolution approaches such as X-ray crystallography or cryo-EM. Furthermore, functional characterization and evaluation of the ferritin-PCBP axis as a potential therapeutic target may provide new opportunities for developing antimalarial strategies. Collectively, our findings highlight a previously unexplored aspect of iron regulation at the host-parasite interface that could potentially be targeted for antimalarial intervention.

## Materials and methods

### Mosquito rearing and maintenance

*Anopheles stephensi* female mosquitoes were reared as described in a previous study (33). *Aedes aegypti* laboratory colony, established from field-collected specimens obtained in Assam, India, in 2022, was maintained at the ICMR–National Institute of Virology insectary. Mosquitoes were reared at 28 °C and 70% relative humidity under a 12 h light:12 h dark cycle. Larvae were fed a mixture of dog biscuit and dry yeast (6:4, w/w), while adults were provided with 10% sucrose solution ad libitum. Four-day-old females were blood-fed on live chickens for colony propagation.

### Expression and Purification of Recombinant Proteins

The plasmid encoding *As*PCBP (UniProt ID: A0A182YKR7) and *Hs*PCBP2 (UniProt ID: Q15366) in the pET-28a vector was obtained from BIOMATIK, Canada (Table S3). Primers encompassing the KH1-KH2 domains (*As*PCBPΔKH3) were designed and ordered from Eurofins. The truncated construct, *As*PCBPΔKH3 (containing KH1-KH2 domains), was subsequently cloned into pET-28a. All the recombinant PCBPs were expressed in E. coli BL21(DE3) cells by growing the cultures overnight at 37 °C in presence of antibiotics (kanamycin,50μg/ml) and subsequently transferring it to a larger culture medium at a final concentration of 2%. The culture was then grown to an OD_600_ of ~0.6, followed by induction with 0.5 mM IPTG and overnight incubation at 16 °C. The bacterial pellets collected were resuspended in lysis buffer (50 mM Na_2_HPO_4_, 300 mM NaCl, pH 7.5) and purified using Ni-NTA affinity chromatography with imidazole gradient elution in ÄKTA pure™. Recombinant MqFtn and HfFtn were purified as described previously(33).

### Immunofluorescence Assay

Animal-related experiments were performed following standard protocols and with prior approval from the Institutional Animal Ethics Committees of ICMR-NIMR, New Delhi, India (IAEC/NIMR/2019-1/09) and BITS Pilani, Hyderabad, India (BITS-HYD-IAEC-2025-064). Purified *As*PCBP, *Hs*PCBP2, MqFtn and Hftn was used to raise polyclonal antibodies as described previously(48).

Adult females were blood-fed using a thermally regulated membrane feeder to mimic host body temperature. Whole bodies, midguts, and ovaries were collected at 24, 48, and 72 hrs post blood meal. Third-instar larvae were also collected. Samples for confocal microscopy were prepared following previously described protocols (33, 49).

### Tissue collection and RNA extraction

To investigate the expression dynamics of iron homeostasis-related genes, samples from different developmental stages of *Aedes aegypti* (eggs, L1–L4 larvae, pupae, and adult males and females) were collected in TRIzol reagent after removing excess water. Tissue-specific expression was evaluated in the midgut, ovaries, fat body, and carcass dissected from 4–5-day-old sugar-fed females. Blood meal-induced changes in gene expression were assessed in midgut and ovary tissues collected at 12, 24, 48, and 72 h post-blood meal. Total RNA was isolated using the TRIzol method and quantified using a NanoDrop 1000 spectrophotometer (Thermo Fisher Scientific).

### cDNA preparation and gene expression analysis

Approximately 1.5 μg of total RNA was reverse-transcribed to first-strand cDNA using the PrimeScript™ 1st Strand cDNA Synthesis Kit (Takara Bio, Kusatsu, Japan) according to the manufacturer’s instructions. Quantitative real-time PCR (qRT-PCR) was performed using SYBR Green qPCR Master Mix (Takara Bio, Kusatsu, Japan) on a CFX96™ Real-Time PCR Detection System (Bio-Rad, USA). The amplification protocol consisted of an initial denaturation step at 95°C for 5 min, followed by 40 cycles of denaturation at 95°C for 10 s, annealing at 56°C for 15 s, and extension at 72°C for 22 s. Fluorescence signals were acquired at the end of each extension step. Amplification specificity was verified by melt-curve analysis using a dissociation program of 95°C for 15 s, 65°C for 5 s, and 95°C for 50 s. All experiments were performed with three independent biological replicates. The ribosomal protein gene *Rps7* was used as the internal reference gene for normalization (Table S4). Relative transcript abundance was calculated using the 2^™ΔΔCt method. Expression profiles were visualized using OriginPro 8.1 software (OriginLab Corporation, USA). Data are presented as mean ± SEM of three biological replicates. Differences between groups were analysed using a two-tailed Student’s t-test, and significance levels are denoted as P ≤ 0.05 (^*^), P ≤ 0.005 (^**^) and P ≤ 0.0005 (^***^)

### dsRNA-mediated gene silencing

To investigate the functional role of PCBP in *Aedes aegypti*, gene silencing was performed using dsRNA-mediated RNA interference (RNAi). Double-stranded RNA (dsRNA) targeting *Aedes aegypti* PCBP homolog was synthesized by *in vitro* transcription using the MEGAscript™ RNAi Kit (Thermo Fisher Scientific; Cat. No. AM1626) according to the manufacturer’s instructions (Table S4). Approximately 100 nL of purified dsRNA (~400 bp; ~3 μg μL^−1^) was injected into the thorax of cold-anesthetized, 2-day-old female mosquitoes using a Nanoject III microinjector (Drummond Scientific, CA, USA; Cat. No. 3-000-207). As a negative control, age-matched females were injected with dsRNA targeting the bacterial *lacZ* gene (Table S4). To evaluate the impact of dietary iron following PCBP silencing, mosquitoes were offered a blood meal 48 h post-dsRNA injection. Fully engorged females were separated and maintained under standard insectary conditions. Post-blood-feeding survival and reproductive output, measured as the number of developed oocytes, were subsequently recorded and compared between treatment groups.

### Immunodetection of Protein-Protein Interaction

Interaction between *As*PCBP/*As*PCBPΔKH3 and MqFtn was assessed using Far-Western blotting as described in a previous study(50). Following transfer of the bait protein (MqFtn) onto a nitrocellulose membrane/96 well ELISA plate, the membrane was blocked for 1 hour with 5% skim milk. The membrane was then incubated with 0.2 mg/mL of the prey protein (*As*PCBP or *As*PCBPΔKH3) for 2 hours, while a control membrane was incubated with 0.2 mg/mL bovine serum albumin (BSA). This was followed by three washes with PBST, 10 minutes each. All the membranes were subsequently incubated with anti-*As*PCBP polyclonal antibody (1:2000), followed by three PBST washes (10 minutes each), and then incubated with anti-mouse secondary antibody (1:5000). After three additional PBST washes (10 minutes each), the signal was developed using an enhanced chemiluminescence (ECL) substrate (Thermo) and TMB substrate (SRL) for ELISA. All experiments were performed in triplicate.

### Microscale thermophoresis

Quantification of the binding affinity between *As*PCBP/*As*PCBPΔKH3 and MqFtn/Hftn was performed using a Monolith X (MM-283) instrument (NanoTemper Technologies, Munich, Germany). Proteins (*As*PCBP, *As*PCBPΔKH3, MqFtn, and Hftn) at a concentration of 200 nM were labeled by incubation with His-tag labeling dye (MO-L018 His-Tag Labeling Kit RED-tris-NTA, 2nd Generation) for 30 minutes. Labeled MqFtn and Hftn were mixed with *As*PCBP (46.8 µM) or *As*PCBPΔKH3 (21.8 µM) in a 1:1 ratio, followed by 16 consecutive serial dilutions and incubation for 1 hour. Samples were centrifuged at 11,000 × g for 10 minutes and loaded into Monolith premium capillaries (MO-K025) before measurement on the Monolith X instrument.

To quantify the binding affinity between *As*PCBP/*As*PCBPΔKH3 and ferrous iron, the same experimental conditions were applied while using FeSO_4_ (5 mM) as the ligand. Binding data were plotted as fraction bound to generate dose-response curves. All experiments were performed in duplicates.

### Ferritin iron loading assay

To assess iron delivery by *As*PCBP/*As*PCBPΔKH3 to MqFtn, an iron oxidation assay was performed. Purified *As*PCBP or *As*PCBPΔKH3 (6 µM) was mixed with MqFtn (3 µM) at a 2:1 molar ratio. Absorbance was recorded immediately at 310 nm for 20 minutes at 30-second intervals using a UV-Vis spectrophotometer (V-650, JASCO).

To evaluate the effect of iron loading or depletion on the reaction kinetics, *As*PCBP/*As*PCBPΔKH3 was pretreated with either 100 µM FeSO_4_ or 100 µM EDTA for 2 hours, followed by overnight dialysis (1:1000) at 4 °C. The treated samples were subsequently used for the iron oxidation assay under the same conditions described above. All experiments were performed in triplicate. The data was analysed using GraphPad Prism 9 software (La Jolla, CA, USA).

### In silico Analysis

Sequences for vector PCBPs were obtained from VectorBase (https://vectorbase.org/vectorbase/app/), Fungi PCBP was obtained from FungiDB (https://fungidb.org/fungidb/app), while *Hs*PCBP sequence were retrieved from UniProt (https://www.uniprot.org/). The class divergence mapping was done using TimeTree (https://timetree.org/) (51). Multiple sequence alignment was performed using Jalview, and phylogenetic tree construction was carried out using MEGA as described previously (33) (Table S5). The structures of *As*PCBP, *As*PCBPΔKH3, and MqFtn were predicted using trRosetta (https://yanglab.qd.sdu.edu.cn/trRosetta/) and subsequently refined using GalaxyWeb (https://galaxy.seoklab.org/), while the structure of Hftn (PDB ID: 4Y08) was obtained from the Protein Data Bank (https://www.rcsb.org/). Protein–protein docking between PCBPs and ferritin cages was performed using ClusPro 2.0 (https://cluspro.org/home.php), and the docked complexes were visualized in PyMOL, followed by interacting residue analysis using PDBsum Generate (https://www.ebi.ac.uk/thornton-srv/databases/pdbsum/Generate.html).

## Acknowledgments

We acknowledge BITS Hyderabad for providing the necessary research facilities and infrastructure to carry out this work; The BITS New Faculty Seed Grant (NFSG) is gratefully acknowledged (Grant Reference: N4/24/1029). We also thank the ICMR–National Institute of Malaria Research for financial support as well as academic and research infrastructure assistance. The authors are grateful to Council of Scientific & Industrial Research (CSIR) (09/0905(12422)/2021-EMR-I) and Department of Biotechnology (DBT) for funding support toward this research. We thank Mr. Satya, and Mr. V. P. Singh for their assistance with insectary maintenance. We thank Dr. Ryuji Yanase, National Institute of Infectious Diseases, Japan Institute for Health Security, Japan, for his valuable insights and suggestions that helped improve the manuscript. We also acknowledge the use of ChatGPT-5.5 for proofreading and correcting typographical errors.

