## Supporting Information for "A Poly(rC)-Binding Protein Mediates Iron Delivery to Ferritin in *Anopheles stephensi*"

**Author Emails and ORCID details:**

### Index

#### 1. Supporting information figures

**Figure S1.** Co-expression/localization of *As*PCBP and ferritin in *Anopheles stephensi*--------------------------- 3

**Figure S2.** Comparative docking and binding site analysis of PCBP-ferritin complexes------------------------- 4

**Figure S3.** Comparative electrostatic surface potential of PCBP-ferritin complexes------------------------------ 6

**Figure S4.** Binding and functional analysis of PCBP constructs---------------------------------------------------- 7

#### 2.Tables

**Table S1** Predicted iron binding sites of *As*PCBP -------------------------------------------------------------------7

**Table S2** Comparative electrostatic surface potential maps of PCBP domains across species---------------------8

**Table S3** Amino acid sequence of synthetic construct used in the study--------------------------------------------9

**Table S4** List of primers used in the study---------------------------------------------------------------------------9

**Table S5** List of genes used in phylogenetic tree construction-----------------------------------------------------10

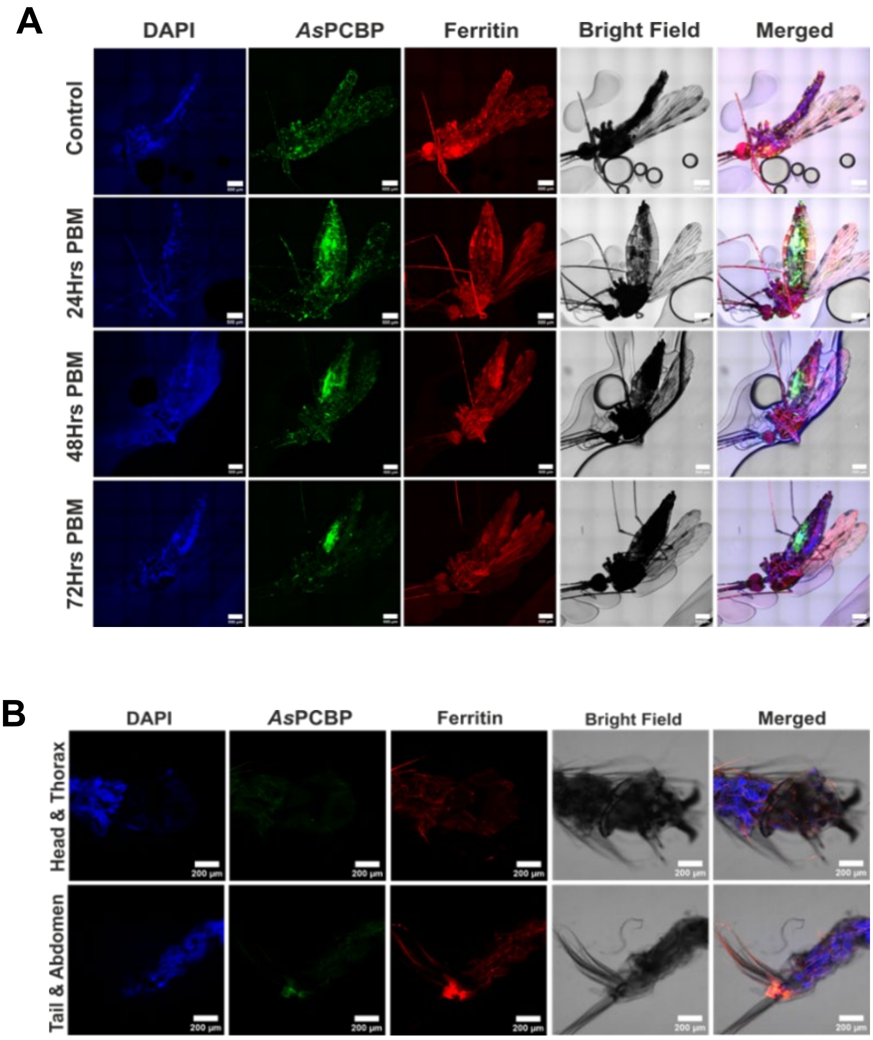


**Figure S1. Co-expression/localization of *As*PCBP and ferritin in *Anopheles stephensi*.** (A) Immunofluorescence analysis of adult female *Anopheles stephensi* at different time points post blood meal (control, 24 hrs, 48 hrs, 72 hrs), and (B) third-instar larvae of *Anopheles stephensi* (bottom panel) showing co-localization of *As*PCBP (Alexa 488-green) and Ferritin (Alexa 647-red) (Scale bar-500µm for adult mosquito and 200µm for larvae).

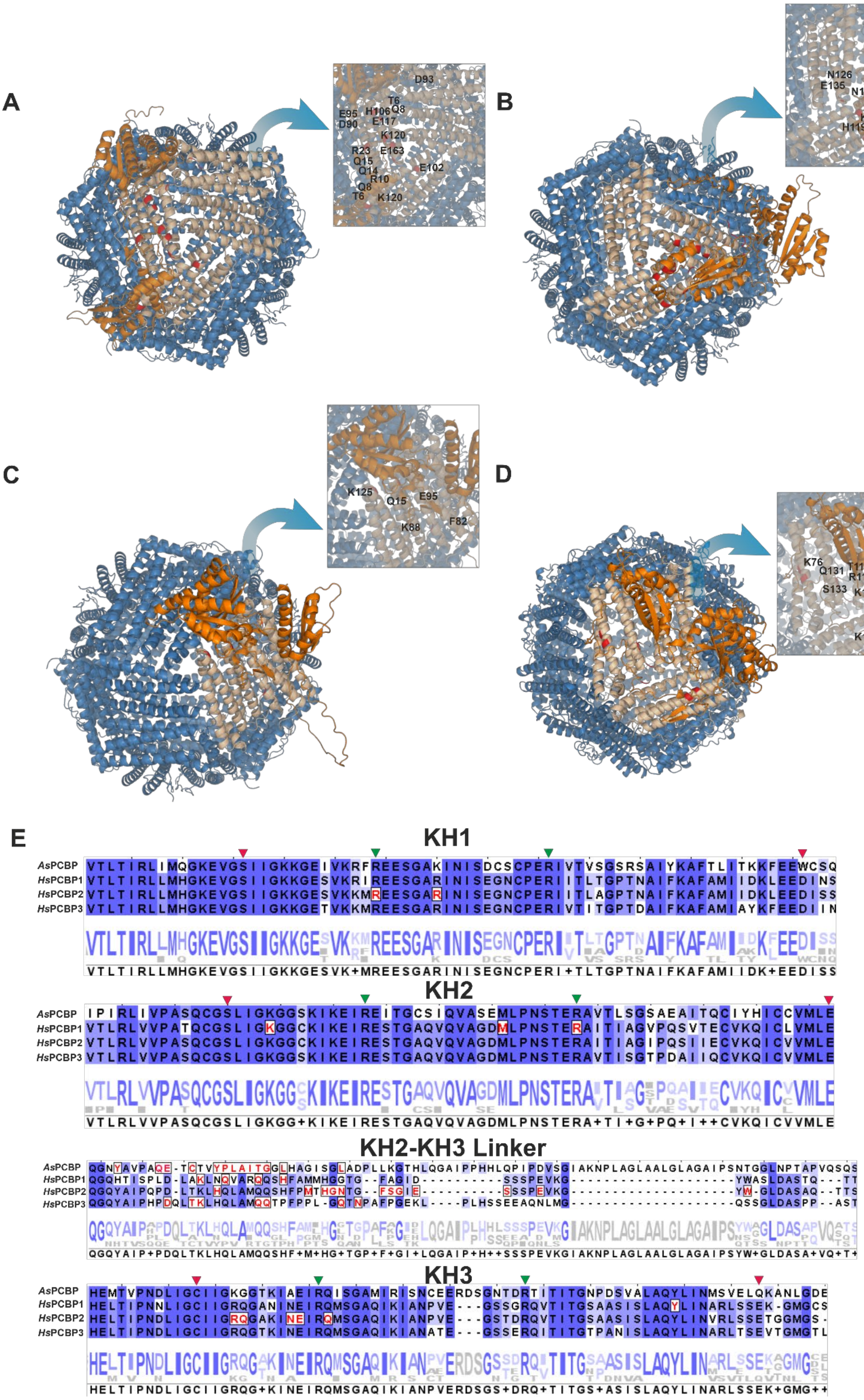
A
D93
T6
Q8
E95
H106
D90
E117
K120
R23
E163
Q15
Q14
E102
R10
Q8
T6
K120
B
D90
D92
E135
N126
K120
D93
E135
N126
E117
Q113
K125
K109
D124
K120
N12
H119
Q15
R23
C
K125
Q15
E95
K88
F82
D
K76
T116
Q131
R114
S133
K110
K204
K140
T201
E
KH1
AsPCBP
HsPCBP1
HsPCBP2
HsPCBP3
KH2
AsPCBP
HsPCBP1
HsPCBP2
HsPCBP3
KH2-KH3 Linker
AsPCBP
HsPCBP1
HsPCBP2
HsPCBP3
KH3
AsPCBP
HsPCBP1
HsPCBP2
HsPCBP3

**Figure S2. Comparative docking and binding site analysis of PCBP-ferritin complexes.** (A-D) Docking models showing the interaction of (A) *Hs*PCBP1 with Hftn, (B) *Hs*PCBP2 with Hftn, (C) *Hs*PCBP3 with Hftn, and (D) *As*PCBP with MqFtn. Insets highlight the interacting residues at the binding interface of each complex. Ferritin is shown in blue, PCBP in orange, the interacting ferritin chain in pale orange, and interacting residues in red.Both *Hs*PCBP1&2 binds to the residue R23, D90 & E117 in Hftn, while Q15 & K125 are common binding site in *Hs*PCBP2&3. Conversely, no common binding residues were found between *Hs*PCBP1&3. (E) Multiple sequence alignment showing the interacting sites in PCBPs (highlighted red) with ferritins. All the PCBPs seems to be interacting with ferritin through their KH2-KH3 linkers. Red arrow-iron binding site, Green arrow-RNA binding site.

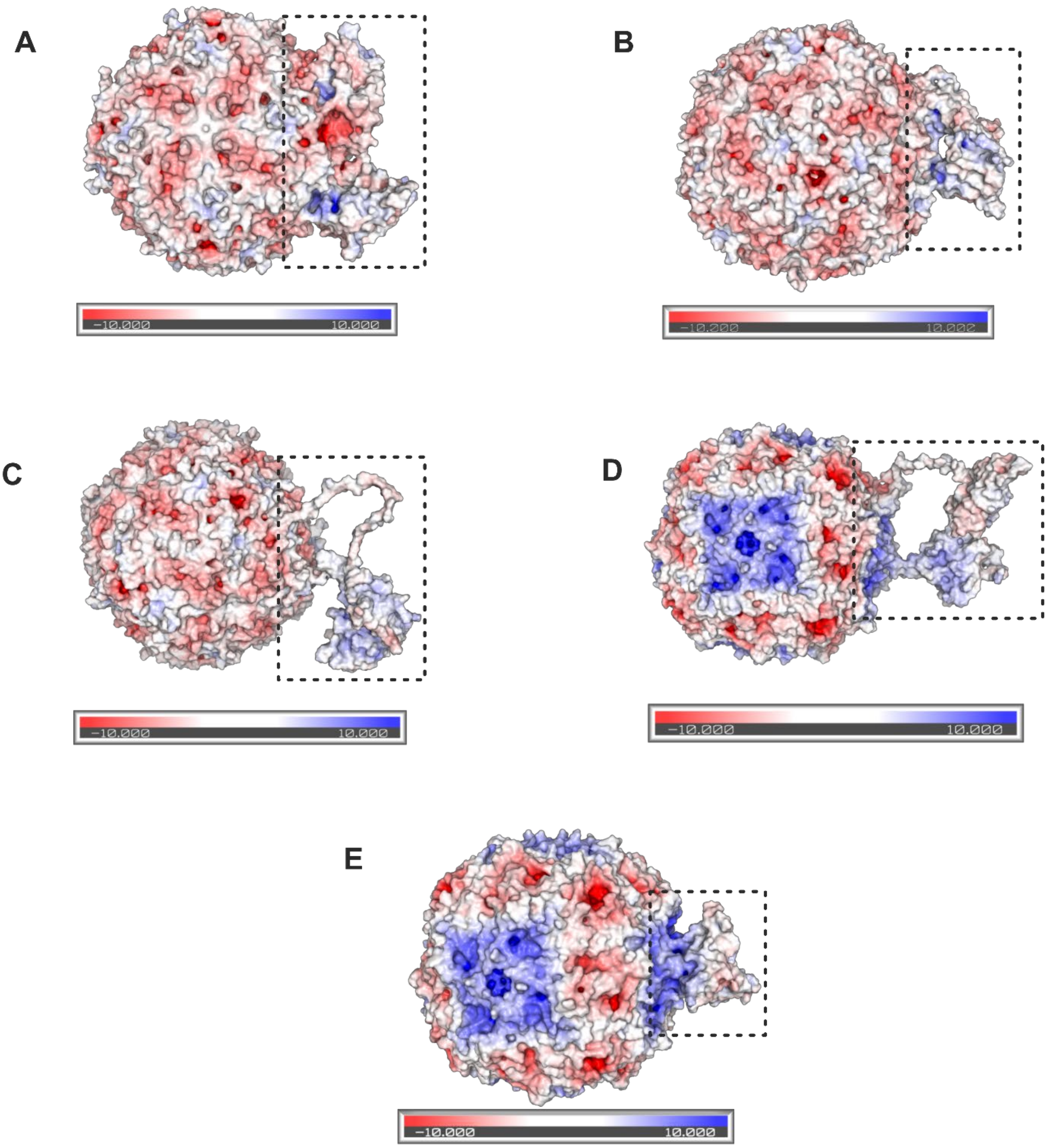


**Figure S3. Comparative electrostatic surface potential of PCBP-ferritin complexes.** (A-E) Electrostatic surface representation of docked complexes showing (A) Hftn with *Hs*PCBP1, (B) Hftn with *Hs*PCBP2, (C) Hftn with *Hs*PCBP3, (D) MqFtn with *As*PCBP, and (E) MqFtn with *As*PCBPΔKH3. The boxed regions highlight the PCBP-binding interface on the ferritin surface. The interaction between PCBP and ferritin seems mostly electronegative, with increase in electropositive interaction seen upon domain truncation. Electrostatic potentials are colored as red (electronegative), white (neutral), and blue (electropositive) and was calculated using the APBS plugin in PyMOL.

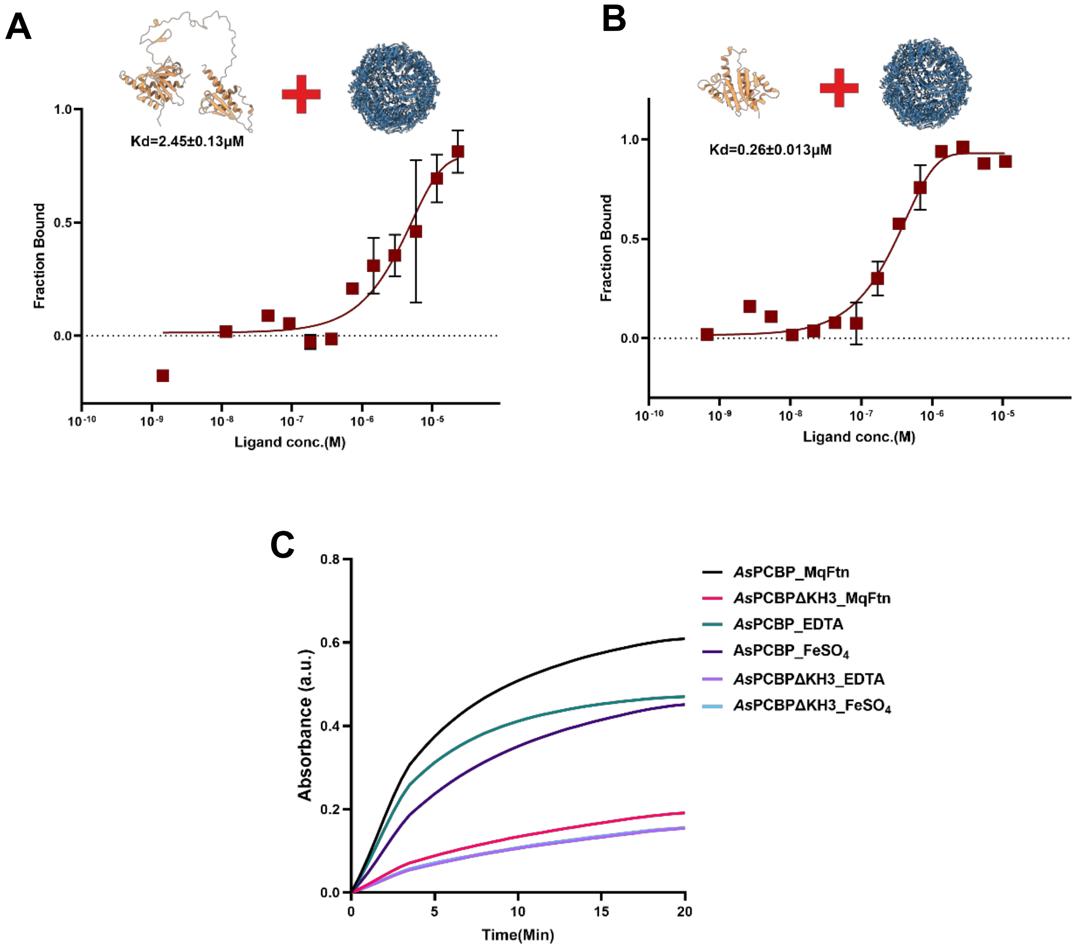


**Figure S4. Binding and functional analysis of PCBP constructs.** (A-B) Microscale thermophoresis (MST) analysis showing binding of (A) *As*PCBP and (B) *As*PCBPΔKH3 to Hftn. Similar to MqFtn, 2-fold increase in binding affinity towards Hftn was observed in the truncated *As*PCBPΔKH3 (C) Iron oxidation kinetics of *As*PCBP and *As*PCBPΔKH3 in the presence of MqFtn following EDTA or $FeSO_4$ treatment. No significant effect seen after EDTA or $FeSO^4$ treatment.

**Table S1.** Predicted iron binding sites of *As*PCBP.

| Predicted iron binding sites (brown) | Predicted iron binding residues | | |
|---|---|---|---|
| | KH1 | KH2 | KH3 |
| 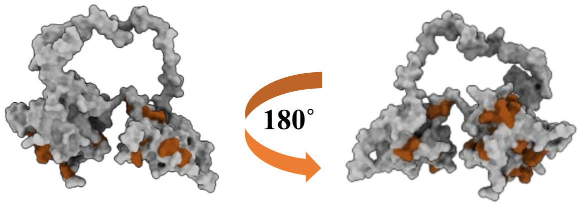  | E42, R46, E50, R73, K85, K86, E88, E89 | K127, K129, E130, R132, E133, Q140, E161, H169 | Q297, H299, E300, E321, Q324, N334, E337, R345, V356, I363, E368 |

*MIB2 server was used for iron binding site prediction.

**Table S2.** Comparative electrostatic surface potential maps of PCBP domains across species.

| | Domain | *Homo sapiens* (PCBP2) | *Anopheles stephensi* |
|---|---|---|---|
| Iron binding site | KH1 | 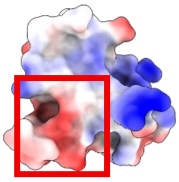 | 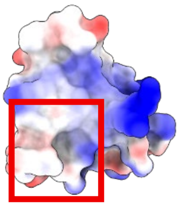 |
| | KH2 | 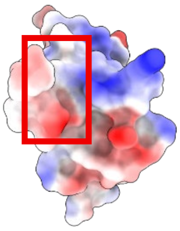 | 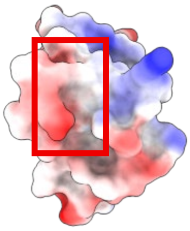 |
| | KH3 | 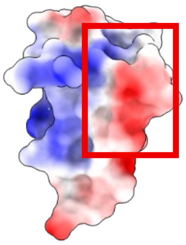 | 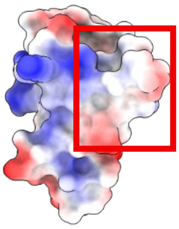 |
| RNA/SsDNA binding site | KH1 | 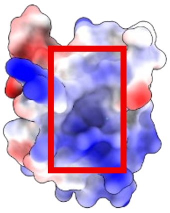 | 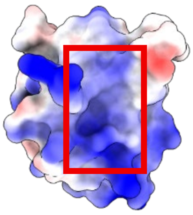 |
| | KH2 | 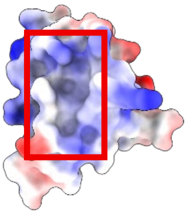 | 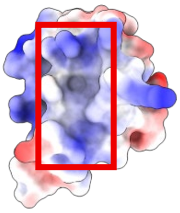 |

| | | | |
|---|---|---|---|
| | **KH3** | 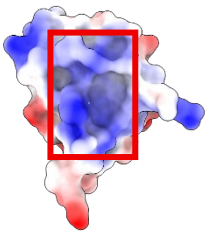 | 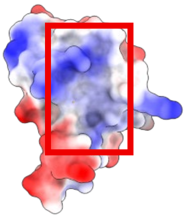 |

**Table S3.** Amino acid sequence of synthetic construct used in the study.

| Protein name (UniProt ID) | Amino acid sequence |
|---|---|
| Hftn (P02794) | MTTASTSQVRQNYHQDSEAAINRQINLELYASYVYLSMSYYFDRDDVALKNFAK YFLHQSHEEREHAEKLMKLQNQRGGRIFLQDIKKPDCDDWESGLNAMECALHL EKNVNQSLLELHKLATDKNDPHLCDFIETHYLNEQVKAIKELGDHVTNLRKMG APESGLAEYLFDKHTLGDSDNES |
| MqFtn (A0A182M0V3) | MQVTDTDSPSGVDEWNYMNRSCSAKLQDQINKEFDAAIFYMQYAAYFAQYKV NLPGFEKFFFEAASEEREHGMKLIEYALMRGQKPIDRNSFSLHFANQVQQPEPEQ GSVALTALRAALAKEQEVTKSIRELIKICEEDHNDYHLVDYLTGEFLEEQHQGQR DLAGKITMLAKLLRTNPKLGEFMFDKQNM |
| *As*PCBP (A0A182YKR7) | MEDTKMVKIGDINLSDDPAVTLTIRLIMQGKEVGSIIGKKGEIVKRFREESGAKINI SDCSCPERIVTVSGSRSAIYKAFTLITKKFEEWCSQFQDQTNTQGKPQIPIRLIVPA SQCGSLIGKGGSKIKEIREITGCSIQVASEMLPNSTERAVTLSGSAEAITQCIYHICC VMLESPPKGATIPYRPKPQVNGPVIVANGQAYTIQGNYAVPAQETCTVYPLAITG GLHAGISGLADPLLKGTHLQGAIPPHHLQPIPDVSGIAKNPLAGLAALGLAGAIPS NTGGLNPTAPVQSQSHEMTVPNDLIGCIIGKGGTKIAEIRQISGAMIRISNCEERDS GNTDRTITITGNPDSVALAQYLINMSVELQKANLGDEAGA |
| *As*PCBPΔKH3 (A0A182YKR7) | MEDTKMVKIGDINLSDDPAVTLTIRLIMQGKEVGSIIGKKGEIVKRFREESGAKINI SDCSCPERIVTVSGSRSAIYKAFTLITKKFEEWCSQFQDQTNTQGKPQIPIRLIVPA SQCGSLIGKGGSKIKEIREITGCSIQVASEMLPNSTERAVTLSGSAEAITQCIYHICC VMLESP |
| *Hs*PCBP2 (Q15366) | MDTGVIEGGLNVTLTIRLLMHGKEVGSIIGKKGESVKKMREESGARINISEGNCP ERIITLAGPTNAIFKAFAMIIDKLEEDISSSMTNSTAASRPPVTLRLVVPASQCGSLI GKGGCKIKEIRESTGAQVQVAGDMLPNSTERAITIAGIPQSIIECVKQICVVMLET LSQSPPKGVTIPYRPKPSSSPVIFAGGQDRYSTGSDSASFPHTTPSMCLNPDLEGPP LEAYTIQGQYAIPQPDLTKLHQLAMQQSHFPMTHGNTGFSGIESSSPEVKGYWGL DASAQTTSHELTIPNDLIGCIIGRQGAKINEIRQMSGAQIKIANPVEGSTDRQVTIT GSAASISLAQYLINVRLSSETGGMGSS |

**Table S4.** List of primers used in the study.

| Name | Forward primer (5'-3') | Reverse Primer (5'-3') |
|---|---|---|
| ***As*PCBPΔKH3 (for cloning)** | CTCGGCTAGCATGGAGGACACCA AGATGGTGAAG | GCGAAAGCTTTAGGGGCTCTCCAG CATCACGCA |

| | | |
|---|---|---|
| **qPCR Ferritin (*Aedes aegypti*)** | CAGCATTGCACCGCCAGATCAA | ACGGGAGCCTTTCCACGCATAA |
| **qPCR Frataxin (*Aedes aegypti*)** | TCCGCCGTTTGCCAAGATGTGT | ACTCCAGCGTGTCGGAACAAAC |
| **qPCR PCBP (*Aedes aegypti*)** | ACTTGGGCTTGCTGGTGCCATT | TTGCCGATGATGCACCCGATCA |
| **SiRNA PCBP (*Aedes aegypti*)** | TAATACGACTCACTATAGGGACGC TCACCATCCGGCTCAT | TAATACGACTCACTATAGGGACGG CACGCTCCGTCGAATTT |
| **SiRNA LacZ** | TAATACGACTCACTATAGGGGAGT CAGTGAGCGAGGAAG | TAATACGACTCACTATAGGGTATCC GCTCACAATTCCACA |
| **qPCR RPS7** | GTGAGCTGGAGAAGAAGTTC | GTCTGCTGGTTCTTGTCC |
| **qPCR Actin** | TATGCCAACACTGTCCTATC | GATTCATCGTACTCCTGCTT |

**Table S5.** List of genes used in phylogenetic tree construction.

| Class | Species | UniProt ID |
|---|---|---|
| **Mammalia** | *Homo sapiens* (*Hs*PCBP1) | Q15365 |
| | *Homo sapiens* (*Hs*PCBP2) | Q15366 |
| | *Homo sapiens* (*Hs*PCBP3) | P57721 |
| | *Homo sapiens* (*Hs*PCBP4) | P57723 |
| | *Myotis myotis* | A0A7J7XJC5 |
| **Aves** | *Gallus gallus* | A0A1D5P893 |
| **Actinopterygii** | *Danio rerio* | A0A8M1PCB6 |
| **Insecta** | *Musca domestica* | A0A1I8MHP1 |
| | *Drosophila melanogaster* | A0A0S0WH03 |
| | *Glossina morsitans* | A0A1B0FGY2 |
| | *Phlebotomus argentipes* | *PARGI1_009222 (VectorBase ID) |
| | *Culicoides sonorensis* | A0A336KRA2 |
| | *Anopheles stephensi* | A0A182YKR7 |
| | *Anopheles gambiae* | A0A499FVL5 |
| | *Anopheles darlingi* | A0A2M4CHN9 |
| | *Aedes aegypti* | Q17LS7 |
| | *Culex quinquefasciatus* | B0X7W6 |
| | *Cimex lectularius* | A0A7E4RKR5 |
| | *Rhodnius prolixus* | A0A4P6D8L1 |

| | | |
|---|---|---|
| **Arachnida** | *Ixodes scapularis* | A0A4D5RG79 |
| | *Rhipicephalus sanguineus* | A0A9D4PRH4 |
| **Sordariomycetes** | *Magnaporthiopsis poae* | A0A0C4DKT0 |
| | *Gaeumannomyces tritici* | J3NRN6 |
| | *Pyricularia oryzae* | G4NBQ1 |
| **Eurotiomycetes** | *Paracoccidioides lutzii* | C1H8C1 |

*UniProt ID not found
